# A Novel Root-Knot Nematode Resistance QTL on Chromosome Vu01 in Cowpea

**DOI:** 10.1101/468215

**Authors:** Arsenio D. Ndeve, Jansen R. P. Santos, William. C. Matthews, Bao L. Huynh, Yi-Ning Guo, Sassoum Lo, Maria Muñoz-Amatriaín, Philip A. Roberts

## Abstract

The root-knot nematode (RKN) species *Meloidogyne incognita* and *M. javanica* cause substantial root system damage and suppress yield of susceptible cowpea cultivars. The narrow-based genetic resistance conferred by the *Rk* gene, present in some commercial cultivars, is not effective against *Rk*-virulent populations found in several cowpea production areas. The dynamics of virulence within RKN populations require a broadening of the genetic base of resistance in elite cowpea cultivars. As part of this goal, F_1_ and F_2_ populations from the cross CB46-Null (susceptible) x FN-2-9-04 (resistant) were phenotyped for *M. javanica* induced root-galling (RG) and egg-mass production (EM) in controlled growth chamber and greenhouse infection assays. In addition, F_2:3_ families of the same cross were phenotyped for RG on field sites infested with *Rk*-avirulent *M. incognita* and *M. javanica*. The response of F_1_ to RG and EM indicated that resistance to RKN in FN-2-9-04 is partially dominant, as supported by the degree of dominance in the F_2_ and F_2:3_ populations. Two QTLs associated with both RG and EM resistance were detected on chromosomes Vu01 and Vu04. The QTL on Vu01 was most effective against aggressive *M. javanica*, whereas both QTLs were effective against avirulent *M. incognita*. Allelism tests with CB46 x FN-2-9-04 progeny indicated that these parents share the same RKN resistance locus on Vu04, but the strong, broad-based resistance in FN-2-9-04 is conferred by the additive effect of the novel resistance QTL on Vu01. This novel resistance in FN-2-9-04 is an important resource for broadening RKN resistance in elite cowpea cultivars.

## INTRODUCTION

Root-knot nematode (RKN) species, particularly *Meloidogyne incognita* and *M. javanica*, cause substantial damage to root systems and suppress yield of susceptible cowpea (*Vigna unguiculata* L. Walp) cultivars by impairing water and nutrient uptake, and the partitioning and translocation of photo-assimilates (Bird and Loveys 1975; McClure 1977; Taylor and Sasser 1978; Williamson and Hussey 1996; Sikora *et al*. 2005). Host-plant resistance is an important strategy to mitigate the impact of nematode infestation (Hall and Frate 1996; Roberts 1992; Ehlers *et al*. 2000; Castagnone-Sereno 2002; National Research Council 2006), both in Africa where access to agronomic inputs including nematicides is limited (Sasser 1980; Luc *et al*. 2005), and in developed agriculture where resistant varieties are the best option economically (Ehlers *et al*. 2000).

Narrow-based resistance to conferred by gene *Rk* has provided protection against RKN in cowpea agricultural systems worldwide (Amosu and Franckowiak 1974; Singh and Reddy 1986; Helms *et al.* 1991; Fery *et al.* 1994; Roberts *et al.* 1995; Roberts *et al.* 1996; Roberts *et al*. 1997; Ehlers and Hall 1997; Ehlers *et al.* 2009). The resistance conferred by gene *Rk* is highly effective against avirulent forms of RKN populations (Roberts *et al*. 1995; Hall and Frate 1996; Roberts *et al*. 1997; Ehlers *et al*. 2000; Roberts *et al*. 2013), but *Rk*-virulent and aggressive forms of common RKN species have been identified (Swanson and Van Gundy 1984; Roberts *et al*. 1995; Hall and Frate 1996; Roberts *et al*. 1997; Petrillo *et al*. 2006). Selection for virulence to *Rk* (Roberts *et al*. 1997; Petrillo and Roberts 2005; Petrillo *et al*. 2006) has prompted efforts to broaden the genetic base of resistance in elite cowpea cultivars (Hall and Frate 1996; Roberts *et al*. 1996; Roberts *et al*. 1997; Ehlers *et al*. 2000; Roberts *et al*. 2013). The threat imposed by virulence in RKN populations led to the discovery of new resistance genes, *Rk^2^* and *rk^3^* to broaden the genetic base of resistance, and advanced breeding materials with one or more of these genes have shown promising performance under RKN infestation (Roberts *et al*. 1996; Roberts *et al*. 1997; Ehlers *et al*. 2000; Ehlers *et al*. 2002). Broad-based genetic resistance can be developed through effective gene pyramiding of independent sets of resistance genes from distinct genetic sources (Ehlers *et al.* 2002).

The RKN resistance currently deployed in many cowpea cultivars is governed by a single dominant gene, *Rk* (Fery *et al.* 1994; Singh and Reddy 1986), but additional resistance genes *Rk^2^*, with a dominant effect, (Roberts *et al*. 1996; Roberts *et al*. 1997; Ehlers *et al*. 2000), and *rk^3^*, with a recessive and additive effect, (Roberts *et al*. 1996; Ehlers *et al*. 2000), have been identified in cowpea backgrounds (Roberts *et al*. 1997; Ehlers *et al*. 2000). The action of gene *Rk^2^* alone is not clearly understood, but in breeding line IT84S-2049 (which also carries gene *Rk*) its additive effect contributes substantially to an enhanced resistance to *Rk*-virulent populations of *M. incognita* and to *M. javanica* compared to gene *Rk* alone (Roberts *et al*. 1996; Roberts *et al*. 1997; Roberts *et al*. 2005). The *rk^3^* locus was characterized as a modifier which improves resistance of cowpea cultivars carrying *Rk* when challenged with *Rk*-virulent RKN isolates (Ehlers *et al*. 2000b) and was bred into cowpea cv. CB27 (Ehlers *et al*. 2000a).

The *Rk* locus has been mapped on chromosome Vu04 (Huynh *et al*. 2016) previous cowpea linkage group 11 of the cowpea consensus genetic map (Lucas *et al*. 2011; Muñoz-Amatriaín *et al*. 2017). This genomic region and flanking markers associated with RKN resistance within this region are important resources for introgressing this resistance into elite cowpea cultivars. Also, markers flanking the resistance in this genomic region can be utilized as a reference to decipher the genetic relationship between the resistance conferred by gene *Rk* and potential novel sources of resistance to RKN.

A broad-based resistance to RKN has been identified through a series of field, greenhouse and seedling growth pouch tests in a cowpea accession FN-2-9-04 from Mozambique (Ndeve *et al*. 2018). This accession carries higher levels of resistance to avirulent *M. incognita* and *M. javanica* than that conferred by the *Rk* gene alone. The performance of FN-2-9-04 under *M. javanica* infestation was contrasted to cowpea breeding lines and cowpea cultivars carrying sets of RKN resistance genes, including *RkRk/Rk^2^Rk^2^*, *RkRk/rk^3^rk^3^*, *RkRk/Rk^2^Rk^2^/gg* and IT84S-2049 which indicated that the RKN resistance in accession FN-2-9-04 is unique. Therefore, to characterize the resistance in FN-2-9-04, genetic analyses were conducted to determine its genomic architecture and localization through genetic linkage analysis and QTL mapping.

### Data Availability

All F_2_ and F_2:3_ populations and root-knot nematode isolates are available upon request. Phenotypic and genotypic date are included in data (D) files 1 - 5. These data files and supplementary tables and figures are available at Figureshare.

## MATERIALS AND METHODS

### Plant materials

Four F_1_, three F_2_ and one F_2:3_ populations (Table 1) were developed under greenhouse conditions at the University of California Riverside (UCR). Accession FN-2-9-04 was crossed with CB46-Null, CB46, Ecute and INIA-41. A single F_1_ seed from each of the crosses CB46-Null x FN-2-9-04, CB46 x FN-2-9-04 and INIA-41 x FN-2-9-04 was grown to derive three independent F_2_ populations, and 150 F_2_ lines of population CB46-Null x FN-2-9-04 were advanced to generate 150 F_2:3_ families (Table 1). Four F_1_ populations (CB46-Null x FN-2-9-04, CB46 x FN-2-9-04, INIA-41 x FN-2-9-04, Ecute x FN-2-9-04) and subsets of their F_2_ populations were phenotyped for root-galling and egg-mass production in greenhouse and seedling growth-pouch screens, respectively, following infection with nematode isolates listed in Table 1. Five to ten seeds per F_1_ population were also screened in each test. The subsets of F_2_ populations and F_2:3_ families (Table 1) also were phenotyped for root-galling in field experiments.

**Table 1.**
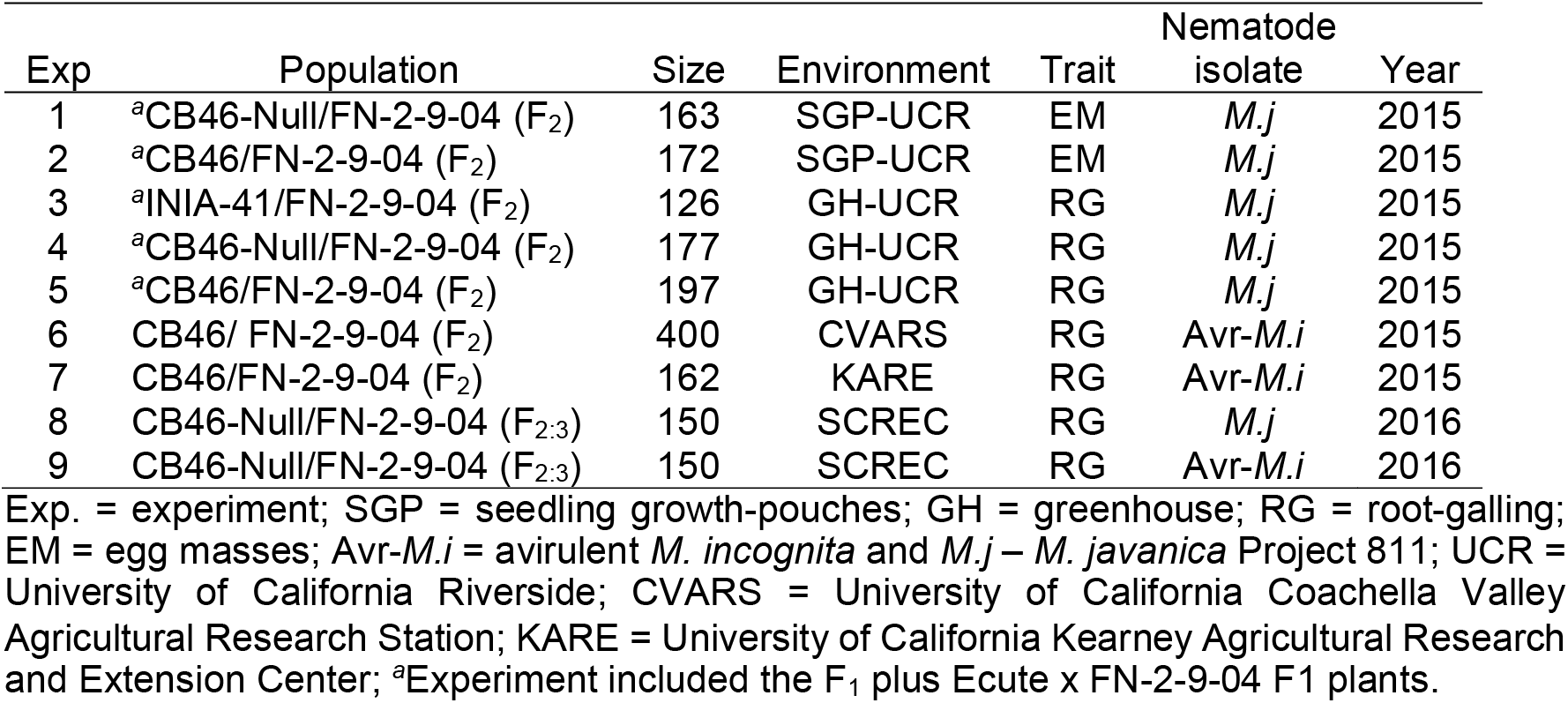
Cowpea populations used for inheritance studies and QTL mapping, their size, phenotyping conditions, target trait, nematode isolate used and year of testing.

| Exp | Population | Size | Environment | Trait | Nematode isolate | Year |
| --- | --- | --- | --- | --- | --- | --- |
| 1 | <sup>a</sup>CB46-Null/FN-2-9-04 (F<sub>2</sub>) | 163 | SGP-UCR | EM | *M.j* | 2015 |
| 2 | <sup>a</sup>CB46/FN-2-9-04 (F<sub>2</sub>) | 172 | SGP-UCR | EM | *M.j* | 2015 |
| 3 | <sup>a</sup>INIA-41/FN-2-9-04 (F<sub>2</sub>) | 126 | GH-UCR | RG | *M.j* | 2015 |
| 4 | <sup>a</sup>CB46-Null/FN-2-9-04 (F<sub>2</sub>) | 177 | GH-UCR | RG | *M.j* | 2015 |
| 5 | <sup>a</sup>CB46/FN-2-9-04 (F<sub>2</sub>) | 197 | GH-UCR | RG | *M.j* | 2015 |
| 6 | CB46/ FN-2-9-04 (F<sub>2</sub>) | 400 | CVARS | RG | Avr-*M.i* | 2015 |
| 7 | CB46/FN-2-9-04 (F<sub>2</sub>) | 162 | KARE | RG | Avr-*M.i* | 2015 |
| 8 | CB46-Null/FN-2-9-04 (F<sub>2:3</sub>) | 150 | SCREC | RG | *M.j* | 2016 |
| 9 | CB46-Null/FN-2-9-04 (F<sub>2:3</sub>) | 150 | SCREC | RG | Avr-*M.i* | 2016 |

Exp. = experiment; SGP = seedling growth-pouches; GH = greenhouse; RG = root-galling; EM = egg masses; Avr-*M.i* = avirulent *M. incognita* and *M.j* – *M. javanica* Project 811; UCR = University of California Riverside; CVARS = University of California Coachella Valley
Agricultural Research Station; KARE = University of California Kearney Agricultural Research and Extension Center; <sup>a</sup>Experiment included the F<sub>1</sub> plus Ecute x FN-2-9-04 F<sub>1</sub> plants.

CB46 is a California blackeye cultivar carrying gene *Rk* (Helms *et al*., 1991), and the CB46-Null genotype is a near-isogenic breeding line (NIL) derived from CB46. This breeding line has the CB46 background, but it is susceptible (minus *Rk* via backcrossing) (Huynh *et al*., 2016). Ecute and INIA-41 are landraces and FN-2-9-04 is an accession from Mozambique. FN-2-9-04 is resistant to both the avirulent *M. incognita* isolates and *M. javanica* isolate used in this study, whereas CB46-Null, CB46, Ecute and INIA-41 are all susceptible to *M. javanica*. In addition, CB46-Null and Ecute are susceptible to the avirulent *M. incognita* isolates (Beltran and Project 77), whereas INIA-41 is resistant.

### Root-knot nematode isolates

Four RKN isolates were used to phenotype plant materials for response to infection. Three *M. incognita* isolates, Beltran, Project 77 and an equivalent isolate indigenous to CVARS are avirulent to the *Rk* gene, with little or no galling and EM production on root systems of plants carrying gene *Rk* (Roberts *et al.,* 1995; Roberts *et al.,* 1996; Roberts *et al.,* 1997), whereas *M. javanica* isolate Project 811 is an aggressive isolate due to its enhanced parasitic ability (Ehlers *et al*., 2000; Ehlers *et al*., 2009), inducing galling and reproducing successfully on roots of plants carrying *Rk* (Thomason and Mckinney, 1960; Roberts *et al.,* 1997; Ehlers *et al*., 2009).

### Resistance phenotyping: egg-mass production

The F_1_ and F_2_ populations (Table 1) plus parental genotypes were phenotyped for *M. javanica* EM production in seedling growth-pouches according to Ehlers *et al*. 2000 and Atamian *et al*., 2012. Briefly, a single seed of each F_1_ and F_2_ was planted per plastic pouch, and the plants were grown in a controlled environment chamber with day/night temperatures set at 28/22 ^o^C under 16 h day-length. Plants were inoculated two weeks after germination with 1500 freshly hatched second-stage juveniles (J_2_) of *M. javanica*. Two days after inoculation, plants were supplied daily with fertilizer for 3-5 days using half-strength Hoagland’s solution (Hoagland and Arnon, 1950). Thirty-five days after inoculation, the pouches were irrigated with erioglaucine dye (Sigma Chemical Co., St. Louis, MO, USA) to stain egg-masses, which were counted under 10X magnification.

### Resistance phenotyping: root-galling

Phenotyping for resistance to root-galling was conducted under greenhouse and field conditions in 2015 and 2016 (Table 1). In the greenhouse, the F_1_ and F_2_ populations and parental genotypes phenotyped for response to *M. javanica* egg-mass production in seedling growth-pouches (in growth chamber conditions) were then transplanted into 4L pots containing soil UC-mix 3 and maintained at 28/22 ^o^C day/night temperatures. After 21 days, each plant was inoculated with 10 ml of *M. javanica* egg suspension in water adjusted to 1000 egg/ml. All greenhouse-grown plants were irrigated twice per day by drip-irrigation for about 90 days to allow seed production, and F_2:3_ seeds were collected from each F_2_ plant. After seed collection, the plant tops were cut at 2 – 3 cm above the soil line, and the roots were washed and scored for root-galling response under 10X magnification, using a 0 - 9 gall index (GI) modified from Bridge and Page (1980): 0 = no galls on root system; 1 = very few, small galls and hard to see; 5 = generally large galls can be seen on the root system and the taproot slightly bumped, with bumps of different sizes; 9 = large galls on the root system, and most lateral roots lost.

Field experiments were conducted in 2015 and 2016 at three sites (Table 1). At CVARS and KARE, 400 and 162 CB46 x FN-2-9-04 F_2_ lines, respectively, were phenotyped for root-galling response to avirulent *M. incognita* (isolate Project 77 at KARE and an equivalent to it at CVARS). In 2016 at SCREC parental genotypes, F_2_ and F_2:3_ populations were phenotyped for root-galling response in separate fields infested with avirulent *M. incognita* isolate Beltran or *M. javanica* (Table 3). In both experiments (Exps. 8 and 9), F_2:3_ families with 25 – 30 plants/family were planted in single plots. The *M. javanica* isolate used in the pot and seedling growth-pouch screens was the same isolate used to infest field sites. For both F_2_ and F_2:3_ generations, 25 - 30 seeds were planted on a 1.5 m-long single row plot, and 60 days after plant emergence plant tops were cut at 2 – 3 cm above the soil line, and the root systems dug and evaluated for root-galling using the same root-galling index described for the pot tests (Bridge and Page 1980).

**Table 3.** Best fit segregation ratios (resistant:susceptible) in 119 and 141 F_2_ plants from crosses CB46-Null x FN-2-9-04 and CB46 x FN-2-9-04, respectively, determined using SNP marker loci at the two nematode resistance QTL regions.

| F<sub>2</sub><br>Population | Genotypes<br>(Observed) |  |  | X<sup>2</sup> | P<br>value | Trait | Vu | Isolate |
| --- | --- | --- | --- | --- | --- | --- | --- | --- |
|  | BB + AB | AA | Exp |  |  |  |  |  |
|  | 96 | 23 | 13:3<sup>a</sup> | 0.002 | 0.95-0.99 | RG | 1 |  |
| CB46-NullxFN-2-9-04 | 93 | 26 | 13:3<sup>a</sup> | 0.56 | 0.25-0.50 | RG | 4 | Avr-*M.i* |
| CB46-NullxFN-2-9-04 | 97 | 22 | 13:3<sup>a</sup> | 0.002 | 0.95-0.99 | RG | 1 | *M.j* |
| CB46-NullxFN-2-9-04 | 98 | 21 | 13:3<sup>a</sup> | 0.04 | 0.75-0.90 | EM | 1 | *M.j* |
| CB46xFN-2-9-04 | 111 | 30 | 13:3<sup>a</sup> | 0.44 | 0.50-0.75 | RG | 1 | *M.j* |
| CB46xFN-2-9-04 | 109 | 32 | 13:3<sup>a</sup> | 1.19 | 0.25-0.50 | EM | 1 | *M.j* |

BB = alleles from resistant parent, AB = heterozygous, AA = alleles from susceptible parent; Exp. = expected ratio; RG = root galling, EM = egg masses per root system; Vu = cowpea chromosome naming (Lonardi *et al.*, 2017); Isolate = Nematode isolate; Avr = avirulent *M. incognita* Beltran, *M.j* = *M. javanica*; <sup>a</sup>also fit a 3:1 ratio.

### Inheritance of resistance and allelism test

Segregation for the FN-2-9-04 resistance to root-galling and reproduction by *M. javanica* and root-galling by avirulent *M. incognita* isolates was determined using both phenotypic (root-galling and egg-masses) and genotypic data. In addition, phenotypic data of F_1_, F_2_ and F_2:3_ populations, and SNP marker genotypes of F_2_ populations at mapped QTL regions were processed for goodness-of-fit analysis to determine the genetic model underlying resistance to RKN in FN-2-9-04. Analysis of goodness-of-fit of segregation ratio between resistant-susceptible lines in the F_2_ was performed through marker-trait association analysis using marker genotypes within mapped QTL regions (see Table 2) and phenotypic response of F_2_ and F_2:3_ populations. Each F_2_ line was scored for presence of parental alleles at each locus within the mapped QTL, and scores 2, 1 and 0 were assigned to homozygous favorable allele (BB = resistant parent), heterozygous (AB) and homozygous non-favorable allele (AA = susceptible parent), respectively. The genotype of each F_2_ line, within the QTL region, was determined as the mean score across all marker loci, and it was associated with its RG or EM phenotypic response determined at the F_2_ and F_2:3_ generations. The data for frequency distribution of genotypes (BB, AB and AA) (Table 3) were processed for goodness-of-fit analysis, and the chi-square values were determined following Yates correction for continuity (Little and Hills 1978). The numbers of genetic determinants associated with resistance were estimated using the Castle-Wright (1921) estimator of gene number, 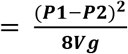, where *n* is the estimated number of genes influencing the trait, *P1* and *P2* are the mean phenotypic values of the parents of the population and *V_g_* is the genetic variance of the trait. To estimate the number of genes governing response to root-galling and egg-mass production, the *V_g_* influencing these traits was derived as the genetic variance in the mapped QTL regions, flanked by known SNP markers.

**Table 2.** Chromosome locations of root-knot nematode (RKN) resistance determinants in cowpea accession FN-2-9-04, mapped using F_2_ and F_2:3_ populations of the cross CB46-Null x FN-2-9-04 and the F_2_ population of the cross CB46 x FN-2-9-04.

| Pop | Trait | RKN | Vu | Position | Flanking markers | -log<sub>p</sub> | PVE (%) | A | D/A |
| --- | --- | --- | --- | --- | --- | --- | --- | --- | --- |
|  |  | Avr- | 1 | 34.4 | 2_04038-2_26991 | 5.4 | 33.0 | -1.3 | 0.5 |
| F<sub>2:3</sub> | RG | *M.i* | 4 | 24.7-27.6 | 2_44685-2_10583 | 20 | 73.3 | -2.0 | 0.5 |
|  | RG | *M.j* | 1 | 27.7-42.0 | 2_47796-1_0027 | 20 | 95.1 | -2.3 | 0.3 |
| F<sub>2</sub> | RG | *M.j* | 1 | 30.3-38.7 | 2_32677-2_19840 | 20 | 47.3 | -2.8 | 0.4 |
| F<sub>2</sub><sup>a</sup> | RG | *M.j* | 1 | 19.2-72.9 | 2_53036-2_18359 | 20 | 65.9 | 2.7 | 0.8 |
| F<sub>2</sub> | EM | *M.j* | 1 | 31.5-36.9 | 2_21671-2_07103 | 10.9 | 34.1 | -17.0 | 0.5 |
| F<sub>2</sub><sup>a</sup> | EM | *M.j* | 1 | 47.1-52.1 | 2_21671-2_12209 | 8.8 | 24.7 | -16.4 | 0.4 |

Pop = mapping population; the F<sub>2:3</sub> were phenotyped in the field whereas the F<sub>2</sub> were phenotyped in greenhouse and growth chamber (seedling-growth pouches) screens; RG = root-galling; EM = egg-masses per root system; Avr-*M.i* = avirulent *M. incognita* isolate Beltran; *M.j* = *M. javanica*; <sup>a</sup>mapping population CB46 x FN-2-9-04 phenotyped for RG and EM; Vu = cowpea chromosome pseudomolecule numbering (Lonardi *et al.* 2017); -log<sub>p</sub> =
level of significance of the detected QTL ( $P < 0.05$ ); PVE = percent of total phenotypic variation explained; A = additive effect of favorable alleles from the resistant parent (negative values indicate the extent of average reduction in RG or EM production due to the presence of favorable alleles; D = dominance effect due to substitution of favorable allele; and D/A = degree of dominance.

Broad-sense heritability (*H^2^ = V_g_/V_p_*) of resistance was estimated using two methods, midparent-offspring regression analysis (Fernandez and Miller 1985; Falconer and Mackay 1996) and the phenotypic variation among F_2_ lines and among F_2:3_ families accounted for by 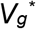 at the QTL regions associated with resistance. The phenotypic variance, *V_ῤ_*, in root-galling or egg-masses attributed to genetic factors, 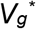, was estimated using SNP marker genotype scores (*V_gs_*) and SNP marker effects (*SNP_eff_*) at the mapped QTL regions plus the observed root-galling or egg-masses phenotypes using the algorithm: 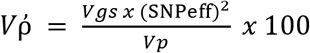. In this algorithm (adapted from Xu 2013), the product *Vgsx* (*SNPeff*)^2^ is the 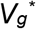 associated with the variation in root-galling or egg-masses phenotypes in tested F_2_ and 2 F_2:3_ populations. To estimate the narrow-sense heritability (*h^2^ = V_a_/V_p_*), the genetic variance 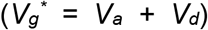 was partitioned into additive and dominance variances, and the *V_a_* component was used to compute the *h^2^* of the trait. Root-galling data of seven F_2_ populations (populations in Table 1 plus their subsets) and parental genotypes were used to perform midparent-offspring regression analysis, and four mapping populations (two F_2_ and two F_2:3_, Exps. 1, 4, 8 and 9, Table 1) were used to derive genetic variances (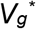) within the QTL regions, influencing the response to galling and egg-mass production. Allelic relationships between the *Rk* locus present in cv. CB46 (Roberts *et al*. 1995; Hall and Frate 1996; Roberts *et al*. 1996; Roberts *et al*. 1997; Ehlers *et al*. 2009; Huynh *et al*. 2016) and the genetic determinants of resistance in FN-2-9-04 were determined using the four F_2_ population sets of CB46 x FN-2-9-04 phenotyped with *M. incognita* isolate Project 77 and *M. javanica* infestation (Table 1).

### Genotyping and QTL mapping

Leaf samples were collected from parents and each of 119 and 137 F_2_ lines of populations CB46-Null x FN-2-9-04 and CB46 x FN-2-9-04, respectively (Exp. 1, 5, Table 1) 30 days after transplanting and dried in plastic ziploc bags containing silica gel packs. Genomic DNA was extracted from dried leaves using Plant DNeasy (Qiagen protocol) and quantified using Quant-iTTM dsDNA Assay Kit and fluorescence measured using a microplate reader. In addition, each F_2_ plant of population CB46-Null x FN-2-9-04 was selfed to generate F_2:3_ seeds for field phenotyping (Table 1). The 119 F_2_ lines are part of the 163 lines tested for egg-mass production (Exp. 1) and transplanted for root-galling assay (Exp. 4, Table 1).

Each DNA sample was assayed for single nucleotide polymorphism (SNP) using the Cowpea iSelect Consortium Array containing 51128 SNPs (Muñoz-Amatriaín *et al*. 2017). The SNP data were filtered for quality as follows: (i) elimination of SNPs with > 20% missing data; (ii) elimination of monomorphic SNPs; (iii) elimination of SNPs with minor allele frequency (MAF) < 0.4 and < 0.3 for populations CB46-Null x FN-2-9-04 and CB46 x FN-2-9-04, respectively; iv) and elimination of duplicated lines. No loci were detected with non-parental alleles.

Linkage-maps of the CB46-Null x FN-2-9-04 and CB46 x FN-2-9-04 F_2_ populations were constructed with MSTmap (Wu *et al*., 2015), and linkage groups were determined at LOD threshold = 10 and marker placement followed the Kosambi mapping function. The options “no mapping size threshold” and “no mapping distance threshold” were fixed at 2 units and 10 cM, respectively. In addition, the no mapping distance threshold option was set at 15 cM and the detection of genotyping errors was not solicited. The linkage groups of the final genetic map were numbered and ordered following the cowpea consensus genetic map order (Muñoz-Amatriaín *et al.* 2017) and the cowpea pseudomolecules (Lonardi *et al*. 2017 in preparation; https://phytozome.jgi.doe.gov/). Also, the cowpea reference genome was used to determine the physical positions of the SNPs and to identify candidate genes on mapped QTLs associated with the traits (Lonardi *et al*. 2017 in preparation; https://phytozome.jgi.doe.gov/). Using physical position, candidate genes were retrieved from the Joint Genome Institute cowpea genome portal (https://phytozome.jgi.doe.gov/pz/portal.html#!info?alias=Org_Vunguiculata_er). QTL mapping was performed using five phenotypic data sets comprising two F_2_ populations of crosses CB46-Null x FN-2-9-04 and CB46 x FN-2-9-04, and two F_2:3_ populations of cross CB46-Null x FN-2-9-04 (Exps. 1, 4, 5, 8 and 9, Table 1). QTL analysis was performed following the mixed-model for QTL mapping described by Xu (2013) using RStudio v1.1.442, and significant QTLs were declared using Bonferroni adjusted threshold value -log (*P*-value) *at P < 0.05*. Reported QTL regions associated with resistance were based on the SNP markers with the most significant threshold values.

### Candidate genes within QTL regions

Single nucleotide polymorphism markers flanking mapped QTL regions on Vu01 and Vu04 were used to determine physical locations of the QTLs and associated candidate genes on the cowpea reference genome v1.0 (Lonardi *et al*. 2017), and a list of gene models and corresponding annotation within each QTL region was generated from the Joint Genome Institute cowpea genome portal (https://phytozome.jgi.doe.gov/pz/portal.html#!info?alias=Org_Vunguiculata_er)

## Results

### Linkage and QTL mapping

The linkage map of the F_2_ population CB46-Null x FN-2-9-04 (n = 119) contained 17208 polymorphic SNP markers distributed on 11 chromosomes and spanned 985.89 cM (Supplementary file S1A). Of the total SNPs, 90.79% (15624 SNPs) were mapped on the cowpea consensus genetic map (Muñoz-Amatriaín *et al*. 2017), while 9.21% (1585 SNPs) were unique to this population, and this portion corresponds to 2.5% of SNPs not mapped to the cowpea pseudomolecules. The linkage map comprised 1392 bins distributed at an average density of 1 bin per 0.71 cM. The linkage map of the F_2_ population CB46 x FN-2-9-04 (n = 137 lines) contained a total of 17903 polymorphic SNPs and spanned 1158.68 cM (Supplementary file S1B). Of these SNPs, 97.6% (17465 SNPs) mapped to the cowpea consensus genetic map, while 9.4% (1675 SNPs) are not part of the cowpea consensus genetic map, and this portion makes 2.4% of the total SNPs not mapped on the cowpea pseudomolecules (Lonardi *et al*. 2017 in preparation; https://phytozome.jgi.doe.gov/).

QTL analysis revealed two major QTLs associated with resistance to root-galling (RG) and egg-mass (EM) production in FN-2-9-04 (Table 2; Figs. 1 and 2); these QTLs were mapped on chromosomes Vu01 and Vu04 of the CB46-Null x FN-2-9-04 population and chromosome Vu04 of the CB46 x FN-2-9-04 population. The QTL region on Vu01 consistently mapped almost within the same genomic location using F_2_ and F_2:3_ populations phenotyped under greenhouse, seedling-growth pouch and field conditions using two RKN isolates (Table 2; Supplementary file S1C).

**Fig. 1.**
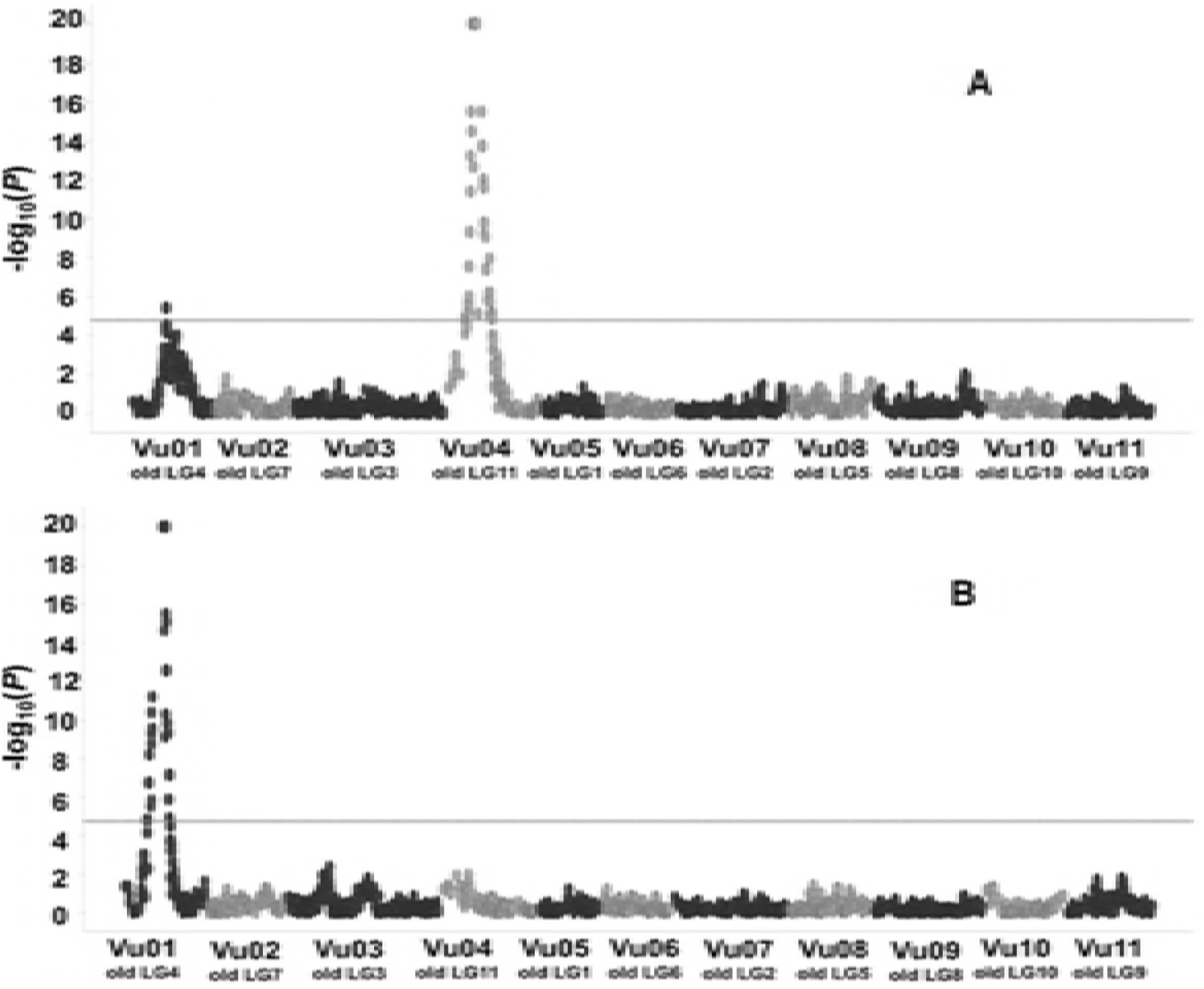
Genomic localization of QTLs associated with resistance to root-galling (RG) by: A, avirulent *M. incognita* and B, aggressive *M. javanica*. The QTLs were detected in the CB46-Null x FN-2-9-04 F_2:3_ population phenotyped for RG undeer field infeststion. Horizontal dashed line represents the Bonferroni thershold of significance at *P* < 0.05 [-log(*p*) = 4.8]. Old LG represents former cowpea linkage group numbering and Vu indicates the new cowpea linkage group numbering based on the cowpea pseudomolecules (lonardi *et al.* 2017).

**Fig. 2.**
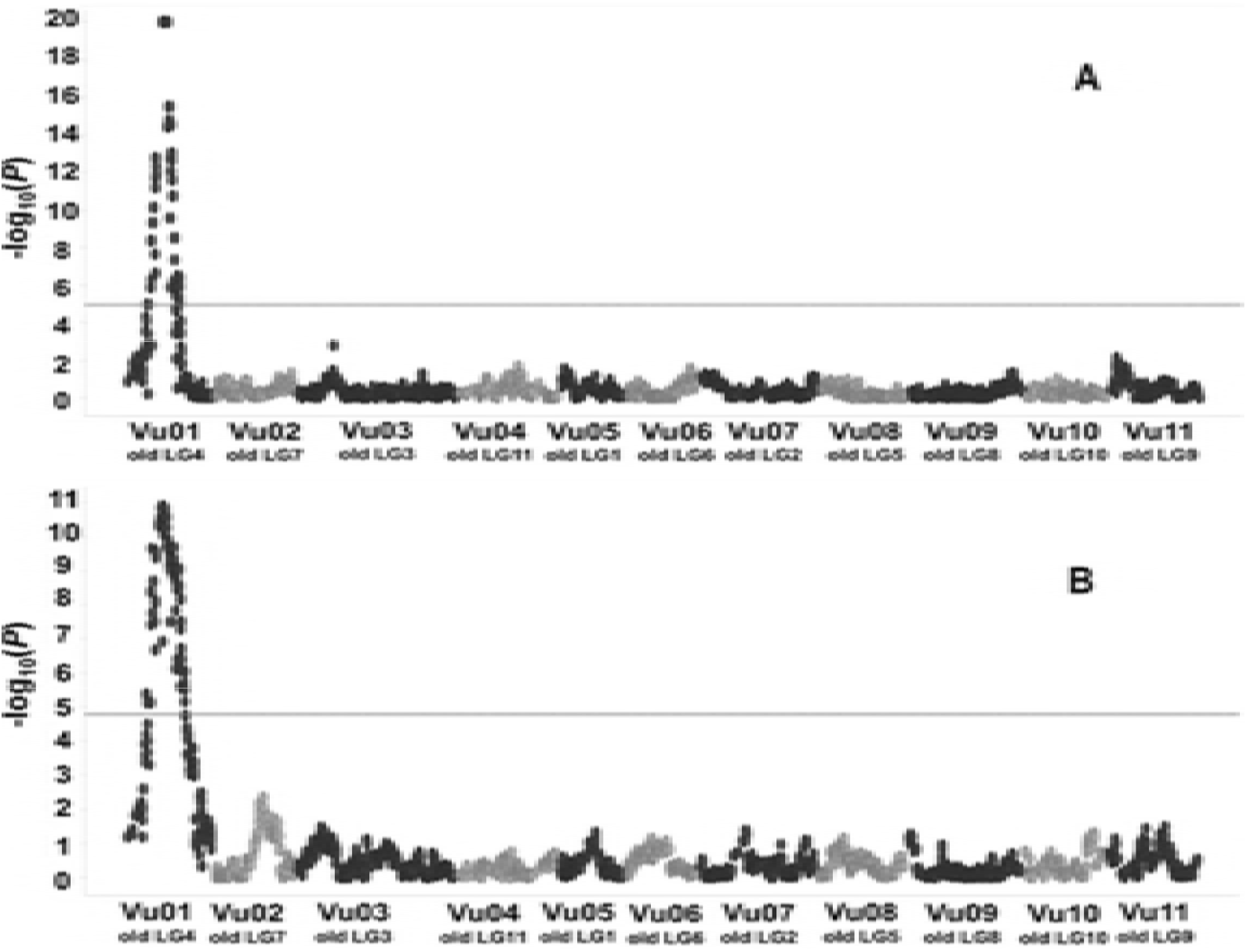
Genomic localization of QTLs associated with resistance to A, root-galling (RG) and B, egg-mass production (EM) by aggressive *M. javanica*. The QTLs were detected in the F_2_ population CB46-Null x FN-2-9-04 phenotyped for RG in the greenhouse and for EM in seedling growth-pouch inoculations, respectively. Horizontal dashed-line represents the Bonferroni thershold of significance at *P* < 0.05 [-log(*p*) (A and B = 4.9 and 4.8, respectively]. Old LG represents former cowpea linkage group numbering and Vu indicates the new cowpea linkage group numbering based on the cowpea pseudomolecules (Lonardi *et al.* 2017).

Two QTLs controlling resistance to RG by avirulent *M. incognita* Beltran were detected and mapped on Vu01 and Vu04 (*P <* 0.05, threshold value -log(*p*) = 4.8) (Fig. 1A) of the CB46-Null x FN-2-9-04 F_2:3_ population. The resistance QTL on Vu01 mapped to position 34.4 cM which spanned 0.1 Mb (28855569 - 28960128 bp) on the cowpea pseudomolecules (Supplementary file S1C) between flanking markers 2_04038 and 2_04039; it accounted for 33% of the total phenotypic variation (*V_p_*) of the RG resistance response and had a likelihood of occurrence expressed by -log_10_(*p*) = 5.4 (Table 2).

The resistance QTL on Vu01 (Fig. 1A) detected under plant infection by avirulent *M. incognita*, exhibited additive and dominance effects of −1.3 and -0.6, respectively, and the degree of dominance, measured as a ratio between dominance and additive effects (D/A), indicated that the resistance in this QTL has partial dominant effect (D/A = 0.5) (Table 2). A second resistance QTL associated with response to the avirulent *M. incognita* was detected on Vu04 (Fig. 1A, Table 2) at chromosome position 24.7 −27.6 cM of the CB46-Null x FN-2-9-04 F_2:3_ population and spanned 2.9 cM which corresponds to approximately 1 Mb (3141521 – 4138458 bp) on the cowpea pseudomolecules (Supplementary file S1C), and it was flanked by SNP markers 2_44685 and 2_10583 (Table 2). This QTL explained 73.3% of the total *V_p_* of the resistance response, and it had an infinite likelihood of occurrence which was represented by -log_10_(*p*) = 20 (Table 2). In addition, the additive (A = −2) and dominance (D = −1) effects of the QTL on Vu04 were slightly higher than those of the QTL on Vu01, but both QTLs showed the same degree of dominance (D/A = 0.5).

On Vu01, an additional genomic region controlling resistance to *M. javanica* RG (Figs. 1B; 2A) and EM production (Fig. 2B) was consistently mapped on the same chromosomal region of the CB46-Null x FN-2-9-04 F_2_ and F_2:3_ populations using RG and EM phenotypic data from field, greenhouse and seedling-growth pouch experiments (Table 2). The *M. javanica* root-galling resistance QTL mapped to positions 30.3 - 38.7 cM and 27.7 - 42.0 cM on Vu01 using F_2_ (greenhouse experiment) and F_2:3_ (Field experiment) populations from the CB46-Null x FN-2-9-04 cross, respectively. These genomic regions spanned 8.4 and 14.3 cM, which correspond to 4.4 (26617356 - 31070755 bp) and 6.2 Mb (25784028 - 31953708 bp) on the cowpea pseudomolecules (Supplementary file S1C) and were flanked by SNP markers 2_32677 - 2_19840 and 2_47796 - 1_0027, respectively (Table 2). In both F_2_ and F_2:3_ populations, the RG resistance QTL was detected with infinite likelihood represented by -log_10_(p) = 20 (Figs. 1B, 2A, Table 2). The percent of total phenotypic variation in RG explained by the QTL effect in the F_2:3_ (PVE = 95.1%) was higher than in the F_2_ (PVE = 47.2%), while the contributions of the additive and dominance effects in the total phenotypic variation in the F_2_ and F_2:3_ were similar (Table 2). Also, the degree of dominance in both generations were comparable, D/A = 0.4 and 0.3, respectively, indicating resistance with partial dominance.

The QTL on Vu01 associated with resistance to *M. javanica* reproduction (EM) mapped to position 31.5-36.9 cM of the CB46-Null x FN-2-9-04 F_2_ population (Fig. 2B; Table 2). This QTL spanned 5.5 cM which corresponds to 2.7Mb (27254299 - 29984745 bp) on the cowpea pseudomolecules (Supplementary file S1C), and it was flanked by SNP markers 2_21671 and 2_07103. This QTL accounted for 34.1% of the total phenotypic variation in EM production with additive and dominance effects of 17.1 and 7.8, respectively; the gene action measured within the same QTL region indicated resistance with partial dominance (D/A = 0.5). Although this QTL was detected with high likelihood, -log_10_(*p*) = 10.9 (critical threshold = 4.8) (Fig. 2B), it was lower than that observed for the RG QTL (Table 2).

QTL mapping using the F_2_ population of CB46 x FN-2-9-04 validated that the genomic region on Vu01 is associated with resistance to *M. javanica* RG (Fig.3; Table 2).

**Fig. 3.**
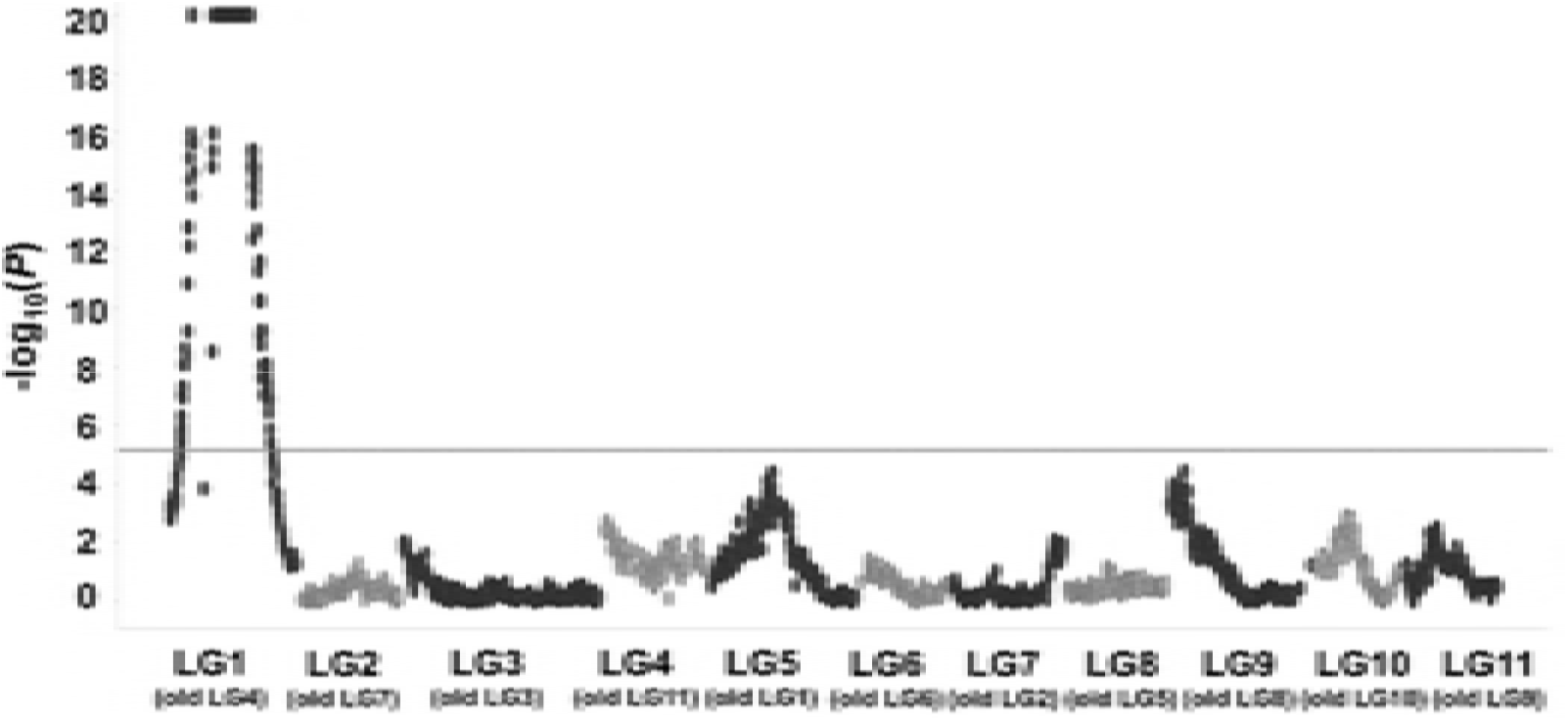
Genomic localization of QTL associated with resistance to root-galling induced by aggressive *M. javanica*. The QTLs was detected in the CB46 x FN-2-9-04 F_2_ population phenotyped for RG in the greenhouse. Horizontal dashed-line represents the Bonferroni thershold of significance at *P* < 0.05 [-log(*p*) = 5.1. Old LG represents former cowpea linkage group numbering and Vu indicates the new cowpea linkage group numbering based on the cowpea pseudomolecules (Lonardi *et al.* 2017).

This Vu01 genomic region was mapped to position 19.2-72.9 cM in the CB46 x FN-2-9-04 F_2_ population, and it spanned 53.7 cM which corresponds to 13.5 Mb (20889089 - 34401992 bp) on the cowpea pseudomolecules with flanking SNP markers 2_53036 - 2_18359 (Table 2; Supplementary file S1C). The QTL on Vu01 explained 65.9% of the total phenotypic variation in *M. javanica* root-galling, and the contribution of the additive and dominance effects were 2.7 and 2.1, respectively. The estimated gene action within this region indicated resistance with partial dominance (D/A = 0.8) (Table 2). This QTL was detected with high likelihood, -log_10_(*p*) = 20 (critical threshold = 5.1) (Fig. 3). In addition, a genomic region associated with resistance to *M. javanica* EM production was mapped on Vu01 of the CB46 x FN-2-9-04 F_2_ at position 46.7 – 53.5 cM, and it spanned 6.8 cM corresponding to 3.2 Mb (27254299 - 30434421 bp) on the cowpea pseudomolecules flanked by SNP markers 2_21671 – 2_12209. This QTL explained 24.7% of the total phenotypic variation in *M. javanica* EM production. (Table 2; Supplementary file S1C).

### Candidate genes within mapped QTL regions

Candidate gene analysis identified a total 316 genes within the genomic region associated with RKN resistance on Vu04 (Supplementary file S2B). Of these, three encode for disease resistance family proteins belonging to leucine rich repeat (LRR) family protein; two genes encode for LRR transmembrane protein kinase; eight encode for disease resistance proteins belonging to toll-interleukin-1-receptor (TIR-NBS_LRR); thirteen genes are putatively considered to also encode for TIR-NBS-LRR class of resistance proteins; one gene encodes for MAP kinase 9; seven genes encode for protein kinase superfamily proteins; three genes encode for receptor-like protein kinase; one gene encodes for pathogenesis-related thaumatin superfamily protein; and two genes encode for TIR-like proteins. Most of these classes of *R* genes were found in adjacent physical positions on the cowpea pseudomolecules.

Within the resistance QTL region on Vu01 a total of 466 genes were identified (Supplementary file S2A). Of these, three encode for LRR family resistance proteins; one gene encodes for TIR-NBS-LRR resistance proteins; eight genes encode for disease resistance-responsive proteins; one gene encodes for hypersensitive-like lesion inducing protein; two genes encode for kinase interaction protein; three encode for LRR protein kinase family protein; one genes encodes for LRR receptor-like protein kinase; three genes encode for LRR and NB-ARC domains-containing disease resistance proteins; fourteen genes encode for NB-ARC domain-containing disease resistance proteins; and four genes encode for protein kinase family proteins.

### Inheritance of resistance in FN-2-9-04

Figures 4A and 4B show the response of four F_1_ populations and their parental genotypes to root-galling (RG) and egg-mass (EM) production, respectively by *M. javanica*. All recurrent parents (Ecute, CB46, INIA-41 and CB46-Null) exhibited susceptible phenotypes for RG and EM, and their mean RG scores and EM scores ranged from 5.8 to 7.7 and 41 to 82, respectively, whereas the resistant parent, FN-2-9-04 had mean RG and EM scores of 0.4 and 4, respectively.

**Fig. 4.**
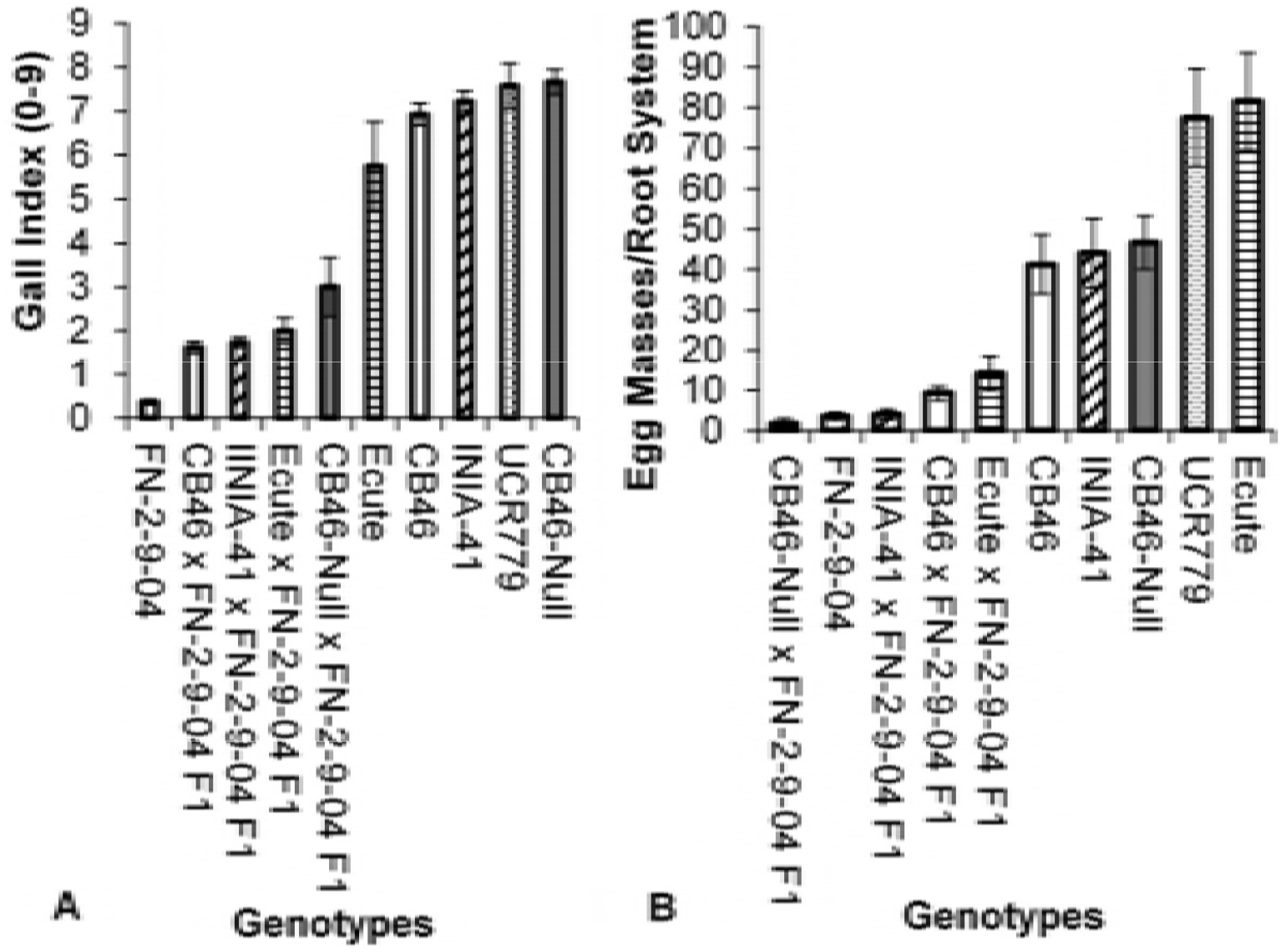
Mean response of F_1_ populations and their parents to: A, root-galling and B, egg-mass production by M. javanica in greenhouse-pot and seeding growth-pouch inoculations, respectively. Bars represent +/- SE.

All F_1_ populations were resistant to *M. javanica* (Fig. 4), with mean RG and EM scores below the mid-parent RG and EM score (GI = 6.9 and EM = 53). The CB46-Null x FN-2-9-04 F_1_ had the highest mean RG (GI = 3) of the four F_1_ populations. The observed differences in RG and EM between the resistant and susceptible parents were significant (*P < 0.05*), but the RG phenotype of the resistant parent was only different from F_1_ populations CB46-Null x FN-2-9-04 and Ecute x FN-2-9-04. The EM phenotypes of the resistant parent and F_1_ were not different. Significant differences among the genotypes were detected at GI = 1.3 and EM = 31.4 (Fig. 4A and 4B).

The segregation of F_2_ (Fig. 5A) and F_2:3_ (Fig. 5B) populations for *M. javanica* RG response appeared to follow a bimodal distribution, skewed toward lower RG phenotype. Also, a bimodal segregation pattern was observed for *M. javanica* EM production in the CB46-Null x FN-2-9-04 and CB46 x FN-2-9-04 F_2_ populations (Fig. 5C). In these same experiments, the average RG observed for parents CB46-Null, CB46, INIA-41 and FN-2-9-09 in greenhouse pots was 7.7, 6.9, 7.2 and 0.4, respectively. In the field experiment (Fig. 5B), RG of 6.7 and 0.1 were observed for parents CB46-Null and FN-2-9-09, respectively, while egg-mass counts per root system equal to 46.7, 45 and 1.8 were observed for parents CB46-Null, CB46 and FN-2-9-09, respectively (seedling-growth pouches).

**Fig. 5.**
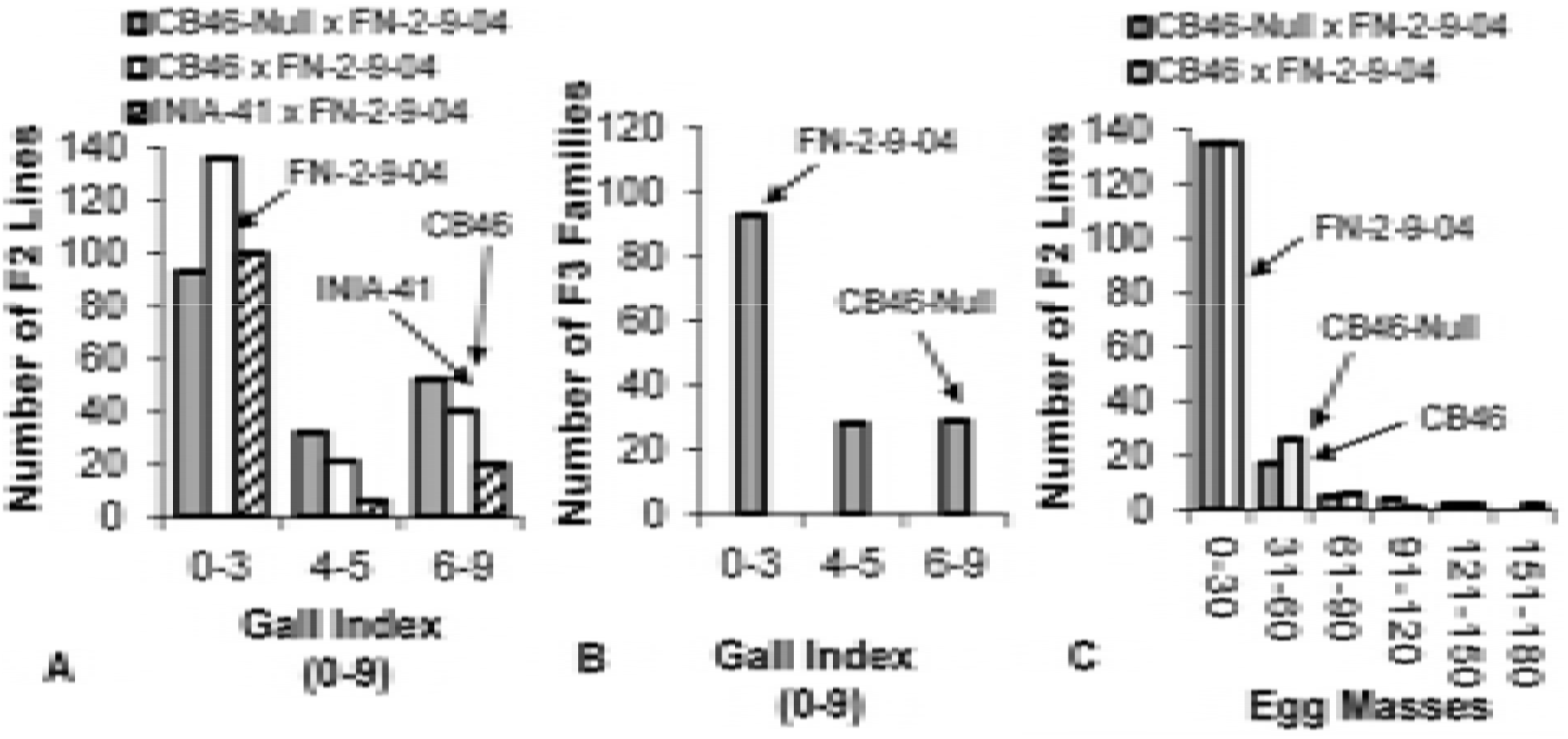
Distribution of root-galling responses in A, F_2_ populations (greenhouse), B, F_2:3_ population CB46-Null x FN-2-9-04 (field), and C, egg-mass production in F_2_ populations CB46-Null x FN-2-9-04 and CB46 x FN-2-9-04 (seeding growth-pouch) under *M. javanica* infestation.

A similar pattern of root-galling distribution was observed in F_2_ (Fig. 6A) and F_2:3_ (Fig. 6B) populations of CB46-Null x FN-2-9-04 under field infestation by avirulent *M. incognita* Beltran. This segregation pattern was consistent across all phenotyping environments (greenhouse, field and seedling growth-pouches) and traits (RG and EM). Egg-mass phenotypes ranged from 0 – 180 (Fig. 5C), and RG across environments and generations ranged from 0 – 9 (Figs. 5 and 6). The resistant parent FN-2-9-04 had consistently lower (*P < 0.*05) RG compared to all susceptible parents. The average *M. incognita* root-galling indices for parents CB46-Null and FN-2-9-04 in the field experiment were 6.4 and 0, respectively.

**Fig. 6.**
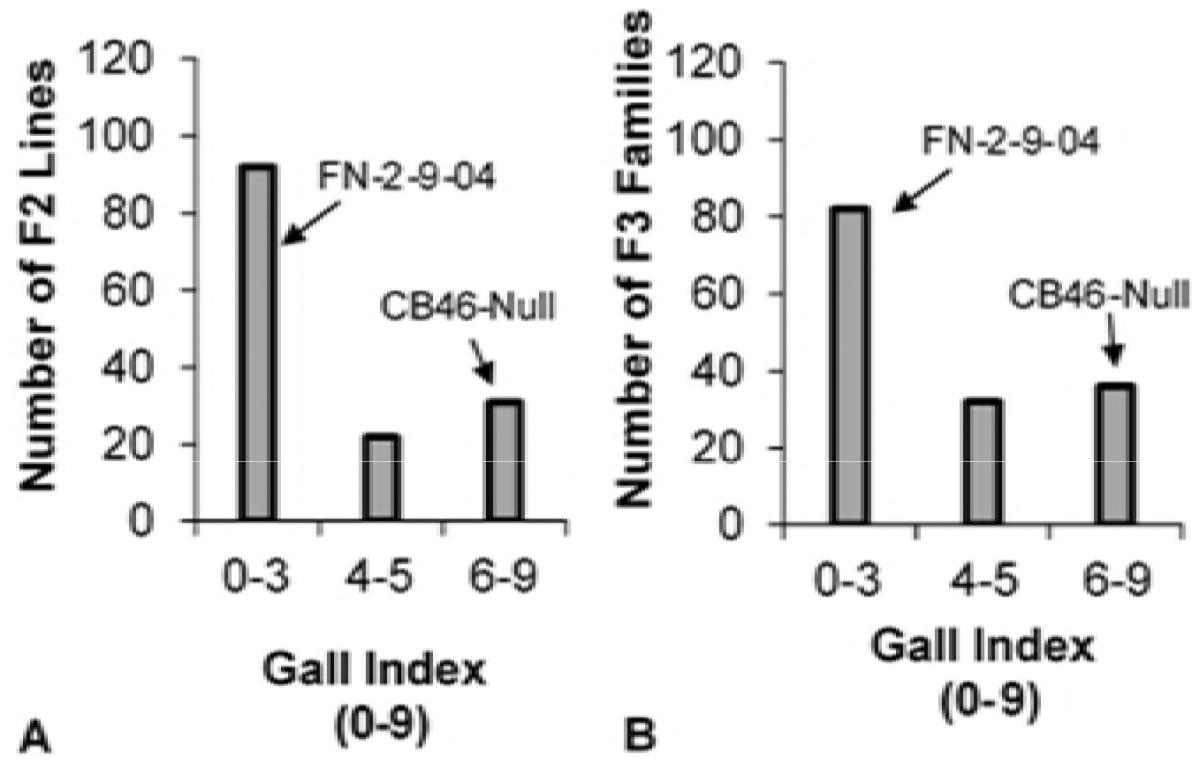
Distribution of root-galling responses in A, F_2_ (A) and F_2:3_ (B) populations of CB46-Null x FN-2-9-04 under field infestation with avirulent *M. incognita* isolate Beltran.

The broad-sense heritability (*H^2^*) of resistance to *M. javanica* root-galling estimated through regression of 7 field phenotyped F_2_ populations to the mean performance of their parents (CB46-Null, CB46, FN-2-9-04 and INIA-41,) was high (*b = 0.76 ± 0.07*, *P = 0.00004*) (Fig. 7), while estimates of *H^2^* for the same trait computed using the genetic variance (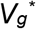) directly derived from the QTL region located on Vu01 were moderate (0.47) and high (0.95) for greenhouse and field phenotyped F_2_ and F_2:3_ populations, respectively. For these populations, the estimates of narrow-sense heritability (*h^2^*) of RG were 0.33 and 0.71, respectively. Egg mass production (EM) response in the F_2_ had low *H^2^* (0.34) (Table 2) and *h^2^* (0.23). The estimated *H^2^* and *h^2^* for resistance to avirulent *M. incognita* RG were 0.33 and 0.23 on Vu01 and 0.73 and 0.49 on Vu04, respectively.

**Fig. 7.**
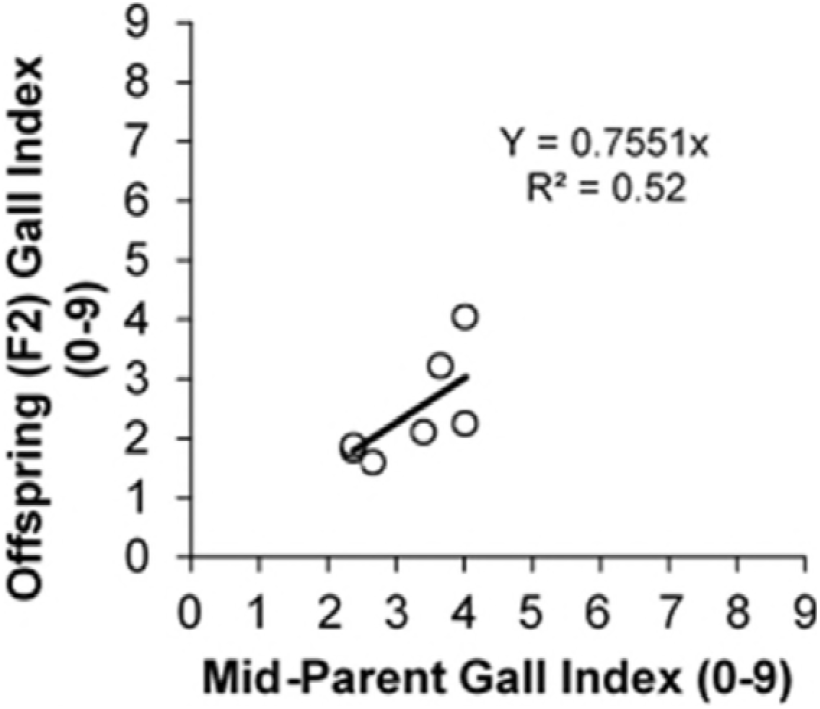
Midparent-offspring regreesion for F_2_ population means regressed on the midparent root-galling values.

Because the *M. javanica* RG and EM resistance QTLs were co-located (Figs. 1B, 2A and 2B), analysis of correlation between RG and EM responses was performed using RG and EM data of F_2_ populations CB46 x FN-2-9-04 and CB46-Null x FN-2-9-04. These traits were highly correlated in both populations, CB46 x FN-2-9-04 and CB46-Null x FN-2-9-04 (*r = 0.78, P = 0.008* and *r = 0.62, P = 0.06*, respectively), although the correlation in the F_2_ population CB46-Null x FN-2-9-04 was not significant (*P = 0.06*) (Fig. 8). The relationship between RG and EM in populations CB46 x FN-2-9-04 and CB46-Null x FN-2-9-04 was explained at 60.3% and 38.1%, respectively, based on the estimated coefficient of determination.

**Fig. 8.**
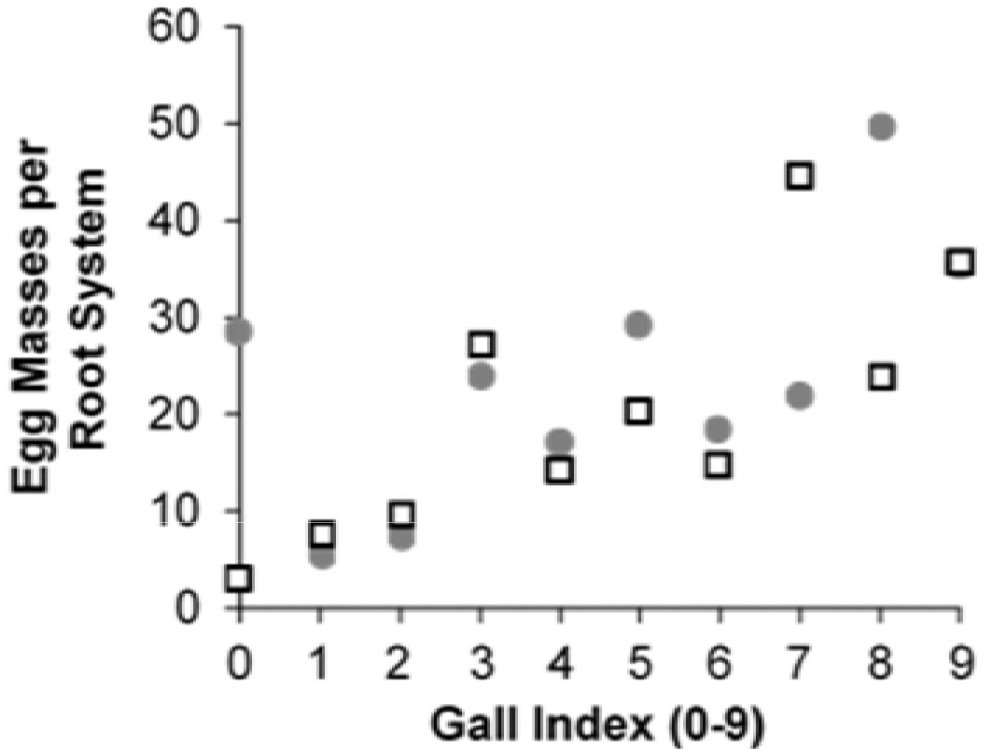
Correlation between *M. javanica* root-galling (greenhouse) and egg-mass production, (seeding growth-pouch) in F_2_ populations • = CB46-Null x FN-2-9-04 (*r = 0.62*) and = □ CB46 x FN-2-9-04 (*r = 0.78*).

The 119 and 137 F_2_ lines of populations CB46-Null x FN-2-9-04 and CB46 x FN-2-9-04, respectively, assayed for 51128 SNP markers segregated for resistance-susceptibility to RG and EM within each mapped QTL, and it fit closely a ratio of 13:3 for phenotypic traits (Table 3). Also, a 3:1 ratio was significant, suggesting that the resistance at both QTL regions is mainly governed by one dominant gene or a combination of genes acting under dominant-recessive interaction. The fit to a 13:3 ratio could also indicate genetic distortion for a single dominant gene.

To validate the genetic models of segregation for resistance-susceptibility to avirulent *M. incognita* and *M. javanica*, gene enumerations were estimated at the mapped QTL regions associated with resistance to RG (Vu01 and Vu04) and EM production (Vu01) following the Castle-Wright (1921) algorithm. The estimates indicated that the resistance to avirulent *M. incognita* RG is under control primarily by 2 and 5 genes residing in QTL regions mapped on Vu04 and Vu01, respectively; whereas, the responses to *M. javanica* RG and EM production mapped on Vu01 are governed mainly by 2 genes each (Supplementary file S3).

Because two QTLs, on Vu01 and Vu04, were associated with resistance to avirulent *M. incognita* RG, analysis of QTLs allele combinations were performed to understand the interaction of both QTLs. Through SNP marker-trait association, the genotype (AA, AB and BB) of each of the 119 F_2_ lines was determined at the QTL regions on Vu01 and Vu04 associated with resistance to avirulent *M. incognita* RG, and each genotype was associated with the average RG phenotypic response of the corresponding F_2:3_. Based on this association, nine QTL combinations (Vu01/Vu04) (Fig. 9) were derived by combining all possible haplotypes on Vu01 and Vu04 contributed from resistant (FN-2-9-04 – favorable allele donor) and susceptible (CB46-Null – non-favorable allele donor) parents.

**Fig. 9.**
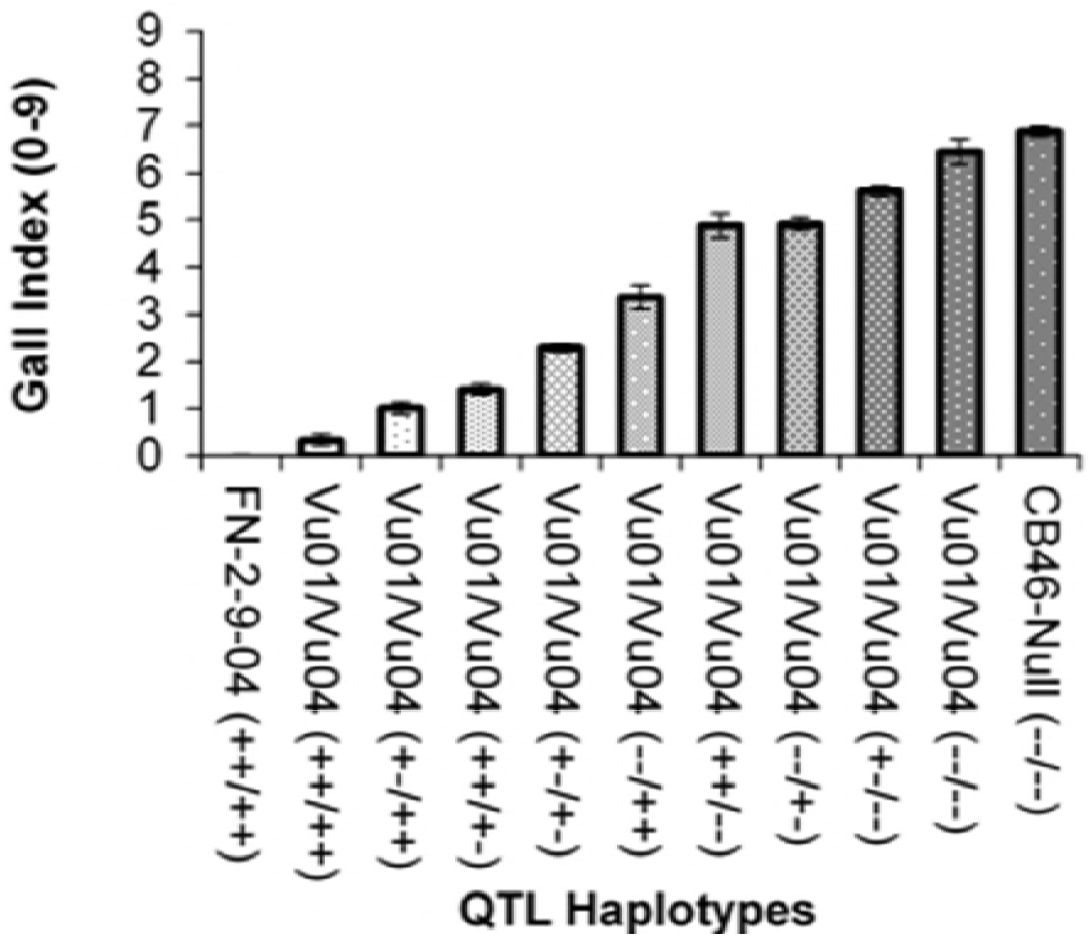
Avirulent *M. incognita* root-galling values for QTL allele combinations for the resistance traits in accession FN-2-9-04 mapped to Vu01 and Vu04 of the cowpea consensus genetic map. The zygosity status within each QTL is indicated by ++, +- and --, representing homozygous favorable, heterozygous and homozygous un-favorable, respectively, in each QTL. Bars are standard errors.

Analysis of variance showed significant effect (*P <* 0.05) of combining QTLs on avirulent *M. incognita* RG response; significant mean differences in RG phenotypes between genotypes carrying combined QTLs were detected at gall index (GI) = 0.88. The resistant parent FN-2-9-04 [Vu01/Vu04(++/++)] did not show any root-galling (Fig. 9), and its response was different (*P <* 0.05) from all genotypes carrying QTL haplotypes with favorable allele dosage different from this parent. Any of the genotypes carrying at least a single favorable allele on at least one of the chromosome regions had less galling than the susceptible parent CB46-Null [Vu01/Vu04(--/--)]. Absence of a single favorable allele in either chromosome predisposed the plants to root-galling, and substantial allele effect was observed for Vu04 [Vu01/Vu04(++/+-)] (Fig. 9). At both loci the favorable alleles must be in the homozygous condition for fully effective *M. incognita* RG resistance.

### Resistance relationship between CB46 and FN-2-9-04

The relationship between the root-galling and nematode reproduction resistance in accession FN-2-9-04 and resistance conferred by the *Rk* gene in CB46 (Huynh *et al*. 2016) was determined through allelism tests using F_2_ populations of CB46 x FN-2-9-04. In addition, analysis of similarity was performed between FN-2-09-04, CB46 and breeding line CB46-Null within the mapped QTL regions to identify putative haplotypes associated with resistance in FN-2-9-04. In 2015 (Table 1), 400 and 162 F_2_ plants plus parents were phenotyped for avirulent *M. incognita* root-galling under field infestation at CVARS and KARE, respectively. At both sites (Fig. 10), all F_2_ plants were resistant with no obvious segregation for root-galling response between plants, indicating that FN-2-9-04 carries a resistance locus allelic to or equivalent to the *Rk* gene found in CB46. The average root-gall indexes for CB46 and FN-2-9-04 were 0.7 and 0.2, respectively.

**Fig. 10.**
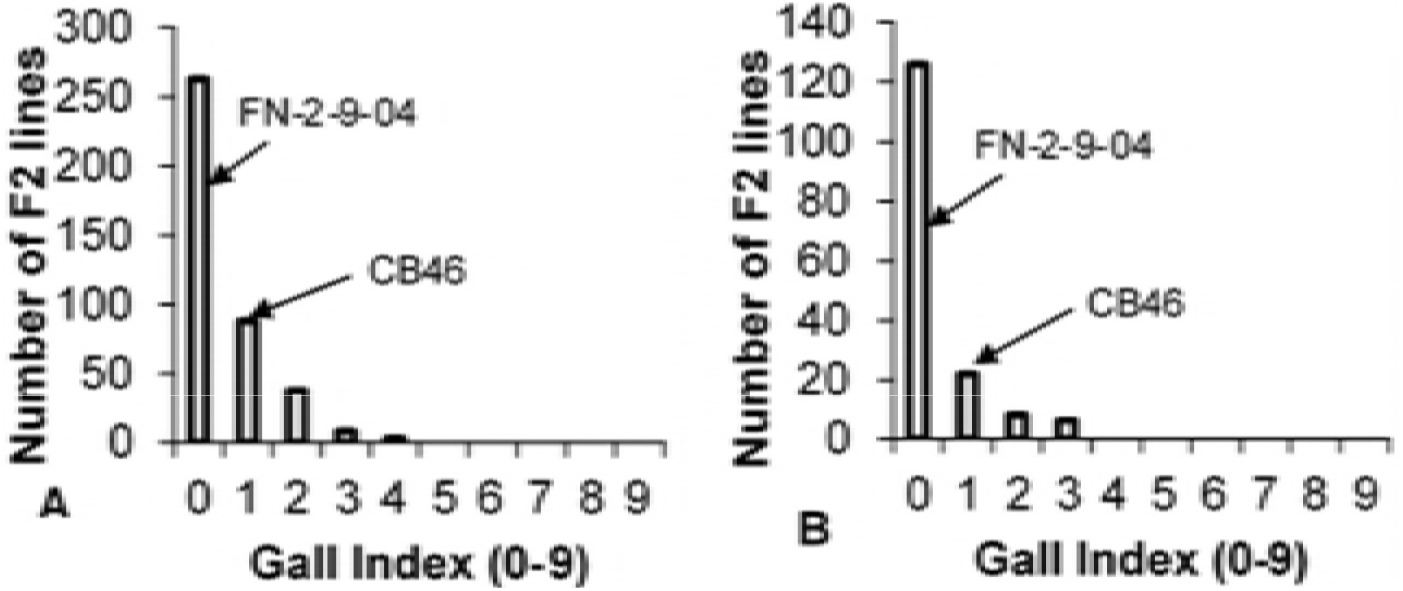
Distribution of root-galling responses in the F_2_ population CB46 x FN-2-9-04 under field infestation by avirulent *M. incognita*(A): Coachella Valley Agricultural Research Station and (B): Kearney Agricultural Research and Extension Center, respectively.

To validate the allelic relationship between resistance determinants conferring resistance to RKN in CB46 and FN-2-9-04, F_2_ population subsets of CB46 x FN-2-9-04 were also phenotyped for resistance to *M. javanica* RG and EM, since these parents exhibited significant differences in *M. javanica* RG and EM production responses (Fig. 4). Using 197 and 172 F_2_ lines for RG and EM phenotyping, respectively (Table 1), segregation occurred for *M. javanica* RG and EM in these F_2_ populations as shown in Figs. 5A and 5C.

Analysis of similarity between FN-2-09-04, CB46 and CB46-Null within the Vu04 genomic region associated with avirulent *M. incognita* RG resistance (Table 2; Fig. 1A) revealed a putative haplotype associated with the resistance (Supplementary file S4). The location of the *Rk* locus on Vu04 identified in CB46 (Huynh *et al*. 2016) overlapped with the resistance region on the same chromosome in FN-2-9-04 within 2.9 cM of the CB46-Null x FN-2-9-04 F_2_ population and within 1.59 cM on the cowpea consensus genetic map (Muñoz-Amatriaín *et al*. 2017), corresponding to approximately 1 Mb on the cowpea pseudomolecules. Within this region, based on SNP marker haplotypes, FN-2-9-04 is 39% identical to CB46 and completely different from CB46-Null (identity = 0%) which is 60% identical to CB46.

Conversely, in the region on Vu01 where an additional resistance QTL was detected in FN-2-09-04 (Table 2; Figs. 1B, 2A, 2B), this resistant parent shares no SNP haplotype similarity with either CB46 or CB46-Null (identity = 0%), whereas CB46 and CB46-Null are 100% identical.

## DISCUSSION

Characterization of the resistance to avirulent *M. incognita* and aggressive *M. javanica* present in cowpea accession FN-2-9-04 from Mozambique revealed that the resistance is determined by two major QTLs which were mapped on chromosomes Vu01 (old LG4) and Vu04 (old LG11) in the CB46-Null x FN-2-9-04 populations and on Vu01 in the CB46 x FN-2-9-04 population.

The QTL mapped on Vu04 overlaps with the previously mapped genomic region which harbors the *Rk* resistance locus (Huynh *et al*. 2016), suggesting that the *Rk* locus is also present in FN-2-9-04. In our previous RKN resistance QTL mapping of *QRk-vu4.1* (old *QRk-vu11.1*) (Huynh *et al*. 2016), this region associated with the *Rk* resistance spanned about 8.35 cM compared to 2.9 cM in this study. This difference in mapping resolution is attributed in part to the current availability of the high-density SNP genotyping platform and high-density cowpea consensus genetic map (Muñoz-Amatriaín *et al*. 2017). If the genomic region harboring the *Rk* locus is a multi-allelic or multi-gene locus, the overlap between *QRk-vu4.1* and the QTL mapped in this study on Vu04 indicates that the resistance alleles are within 2.9 cM interval of the CB46-Null x FN-2-9-04 population corresponding to approximately 1 Mb on the cowpea pseudomolecules. This locus provides effective resistance against avirulent *M. incognita* populations. The resistance to avirulent *M. incognita* present on Vu01 in FN-2-9-04 is confined to 0.1 Mb of the cowpea pseudomolecules, and its relative low contribution to the total phenotypic variation in root-galling response (33%) compared to the resistance in Vu04 (73.3%) supports that the resistance in Vu04 is the main resistance for this nematode although both are required in the FN-2-9-04 background for fully effective resistance. The estimated values of contribution of each resistance QTL to the total phenotypic variance (Vu01 + Vou4; 33% + 73.3%) give a reliable indication of activity of each resistance QTL to the observed root-galling phenotypic response, with the excess in estimation attributed to error.

The resistance to *M. javanica* in FN-2-9-04 consistently mapped to Vu01 using root-galling and egg-mass production phenotypic data from F_2_ and F_2:3_ populations phenotyped under distinct environmental conditions (greenhouse, growth chamber and field). The QTL associated with resistance to *M. javanica* egg-mass production was collocated with the QTL controlling root-galling response, and based on the physical positions, on the cowpea pseudomolecules, of the mapped resistance QTLs, the resistance to *M. javanica* root-galling and egg-mass production are confined within 6.2 Mb. The resistance QTL on Vu01 is distinct from the *Rk* locus (*QRk-vu4.1*, Huynh *et al*. 2016) which was mapped on Vu04, also it is distinct from the recently mapped RKN resistance locus on Vu11 (Old LG9) which also confers resistance to *M. javanica* (Santos *et al*. 2018). Therefore, it represents a novel RKN resistance QTL in cowpea designated here as *QRk-vu1.1*.

The response of four F_1_ populations to root-galling and egg-mass production relative to the resistant parent, and the skewed segregation of these nematode-induced phenotypes in the F_2_ and F_2:3_ populations indicated that these responses are under control by major genes with partial dominance effects, as also indicated by the estimated degrees of dominance (D/A). Resistance to RKN under control by major genes with partial dominance effect has been reported in several studies (Ali *et al*. 2014; Huynh *et al*. 2016).

Analysis of segregation for resistance against *M. javanica* and avirulent *M. incognita* through marker-trait association better fit a 13:3 ratio expected for a genetic control under a single dominant gene plus a recessive gene on both Vu01 and Vu04, also suggesting that the major genes controlling resistance are putatively aided by minor/recessive genes, and collectively in a dominant-recessive interaction to confer substantially stronger, broad-based resistance than that conferred by the *Rk* gene alone. A similar genetic phenomenon of major gene and minor/recessive gene interaction was described in cowpea cultivar CB27, where gene *Rk* acts together with a recessive gene to enhance and broaden root-knot nematode resistance (Ehlers *et al*. 2000). The data also fit a 3:1 ratio expected for a single major gene, and the better fit to the 13:3 of the SNP haplotypes could represent genetic distortion within each locus. However, using the Castle-Wright (1921) algorithm for gene enumeration, the estimates also supported that two genes on Vu01 and two genes on Vu04, may be responsible for the resistance against *M. javanica* and avirulent *M. incognita*, respectively, but the estimates of genes involved in resistance against avirulent *M. incognita* on Vu01 did not support the observed segregation for resistance. The extent of genetic distortion in these regions or multi-allelic effects require further study.

Analysis of candidate genes within QTL regions harboring resistance to root-knot nematode revealed several classes of *R* genes known to be associated with plant disease resistance (Ellis and Jones 2003; Takken and Tameling 2009; Gururania *et al*. 2012); for example, genes encoding for LRR resistance proteins, LRR transmembrane protein kinase, TIR-NBS-LRR resistance proteins, hypersensitive-like lesion inducing protein, and NB-ARC domains-containing disease resistance proteins. The composition and arrangement of these classes of candidate *R* genes identified on QTL regions housed on chromosomes Vu04 and Vu01 were substantially distinct; this phenomenon may explain the specificity and the structure of resistance to root-knot nematode reported in this study. The resistance QTL on Vu04 was specific to *M. incognita* although effective resistance to this nematode was guaranteed by additive effect of the resistance QTL on Vu01, which was specific to *M. javanica*. In both QTL regions, the candidate *R* genes were arranged in tandem. The candidate *R* genes identified in the QTL on Vu04 matched those reported by Santos *et al*. (2018), further supporting that the *Rk* resistance locus is also present in cowpea accession FN-2-9-04. How many of these identified candidate *R* genes are directly involved in determining the RKN resistance phenotypes requires further investigation. The current lack of a functional analysis system in cowpea hampers the determination of which genes are directly involved in the resistance reported here. Therefore, further analysis and testing of function of candidate genes within QTLs associated with resistance to root-knot nematode is a pertinent research goal.

Estimates of heritability of resistance in FN-2-9-04 to avirulent *M. incognita* and aggressive *M. javanica* in the F_2_ generation using greenhouse phenotypic data were lower than those estimated in the F_2:3_ generation using phenotypic data from field experiments. This can be accounted for by the segregation in both populations and because greenhouse phenotyping is less variable compared to field testing. The estimates of narrow-sense heritability of resistance to root-galling induced by both RKN species were in the range 0.23 – 0.71, indicating that the resistance in FN-2-9-04 can be transferred successfully into elite cowpea cultivars to broaden the genetic base of root-knot resistance which currently relies on the *Rk* gene. The resistance response to *M. javanica* reproduction had lower heritability estimates (*H^2^ =* 0.25 and 0.34; *h^2^ =* 0.17 and 0.24) compared to those for *M. javanica* induced root-galling (*H^2^ =* 0.47 - 0.95; *h^2^ =* 0.33 - 0.71), which could be due to egg-mass production data being generally more variable compared to root-galling data. High correlation between root-galling and nematode reproduction responses, and the co-location of resistance QTLs associated with both phenotypes suggests that both traits may be governed by the same genes determining resistance. Similarly, significant correlation between root-galling and reproduction phenotypes in cowpea recombinant inbred populations was reported by Huynh *et al* (2016) for the *Rk* locus on Vu04. In contrast, in lima bean (*Phaseolus lunatus* L.) the responses to root-galling and nematode reproduction were reported to be under control by independent genetic factors (Roberts *et al*. 2008). Since genetic factors explained 38.1 and 60.3 % of the association between root-galling and egg-mass production in this study, these data suggest that although the genomic regions governing both traits are co-located, these traits may be under distinct regulatory mechanisms, or that the resistance to both traits may reside within a multi-allelic locus or tandemly arranged loci.

The heritability of resistance to avirulent *M. incognita* root-galling comprised two components, one on Vu01 (*H^2^ =* 0.33; *h*^2^ *=* 0.23,) and the other on Vu04 (*H^2^ =* 0.73*; h^2^ = 0.49*) indicating that the major locus for this resistance in FN-2-9-04 is housed on Vu04, and it is aided by the additional locus on Vu01 with low resistance heritability. Also, the differential activity between the resistance loci on Vu01 and Vu04 points to specificity of resistance to avirulent *M. incognita* and *M. javanica*. Huynh *et al* (2016) reported that, although the QTL harboring the *Rk* locus had a significant effect on controlling both avirulent *M. incognita* and *M. javanica*, its resistance activity was lower against *M. javanica*. Marker-trait association analysis in the current study indicated that resistances on both Vu01 and Vu04 are required for effective resistance under avirulent *M. incognita* infestation.

The allelism test between CB46 and FN-2-9-04 revealed a lack of resistance segregation in the CB46 x FN-2-9-04 F_2_ population under avirulent *M. incognita* infestation, indicating that both parents carry the same major gene *Rk* locus previously mapped by Huynh *et al* (2016) on Vu04 (old LG11) of the cowpea consensus genetic map (Munoz-Amatriain *et al*. 2017), also supporting that the resistance mapped in this study on Vu04 corresponds to the *Rk* locus. *Rk* was the first identified RKN resistance locus in cowpea, and it has been bred into many commercial cowpea cultivars (Fery and Dukes 1980; Helms *et al*. 1991; Ehlers *et al*. 2009). In contrast, the segregation found in F_2_ population CB46 x FN-2-9-04 for *M. javanica* root-galling and reproduction responses, and the mapping of resistance QTLs for root-galling and egg-mass production confirmed that the heightened and broad-based resistance response in FN-2-9-04 relative to CB46 is conferred by novel resistance determinants located on Vu01.

Flanking markers associated with the mapped genomic regions on Vu01 and Vu04 can be used to assist the introgression of the resistance into elite cowpea cultivars. In particular, the novel resistance detected on Vu01 confers the most effective *M. javanica* resistance known to date in cowpea. The resistance on Vu01 appears to be more specifically effective against aggressive *M. javanica*, while both the Vu01 and Vu04 QTLs have activity against avirulent *M. incognita*, but with the QTL on Vu04 playing the major role in resistance. This was also demonstrated by QTL pyramiding of resistance on Vu01 and Vu04. Thus, both resistance QTLs on Vu01 and Vu04 are responsible for the strong and broad-based resistance observed in FN-2-9-04, which is more effective than the narrow-based resistance provided by the *Rk* gene alone. The mechanism of resistance displayed by this novel broad-based resistance is yet to be determined.

The genetic linkage maps of the F_2_ populations CB46-Null x FN-2-9-04 and CB46 x FN-2-9-04 are additional valuable genetic resources, especially because they are the first cowpea linkage maps constructed using a genotype from the cowpea gene-pool II from southeastern Africa (Huynh *et al*. 2013), and because 9.2% of the 17209 SNP markers on the CB46-Null x FN-2-9-04 map were unique to this population and were not mapped on the most recent version of the cowpea consensus genetic map (Munoz-Amatriain *et al*. 2017).

